# Genetic and structural data on the SARS-CoV-2 Omicron BQ.1 variant reveal its low potential for epidemiological expansion

**DOI:** 10.1101/2022.11.11.516052

**Authors:** Fabio Scarpa, Daria Sanna, Domenico Benvenuto, Alessandra Borsetti, Ilenia Azzena, Marco Casu, Pier Luigi Fiori, Marta Giovanetti, Antonello Maruotti, Giancarlo Ceccarelli, Arnaldo Caruso, Francesca Caccuri, Roberto Cauda, Antonio Cassone, Stefano Pascarella, Massimo Ciccozzi

**Affiliations:** Department of Biomedical Sciences, University of Sassari, Sassari, Italy; Department of Veterinary Medicine, University of Sassari, Sassari, Italy; Unit of Medical Statistics and Molecular Epidemiology, University Campus Bio-Medico of Rome, Rome, Italy; National HIV/AIDS Research Center, Istituto Superiore di Sanità, Rome, Italy; Flavivirus Laboratory, Oswaldo Cruz Institute, FIOCRUZ, Rio de Janeiro, Brazil; Department of Science and Technology for Humans and the Environment, University of Campus Bio-Medico di Roma, 00128 Rome, Italy; Department GEPLI, Libera Università Ss Maria Assunta, 00193 Rome, Italy; Department of Public Health and Infectious Diseases, University Hospital Policlinico Umberto I, Sapienza University of Rome, 00161 Rome, Italy; Department of Molecular and Translational Medicine, Section of Microbiology, University of Brescia, Brescia, Italy; UOC Malattie Infettive, Infectious Disease Department, Fondazione Policlinico Universitario Agostino Gemelli IRCCS, Largo Francesco Vito 1, 00168, Rome, Italy; Universita degli Studi di Siena-Sede di Arezzo, 52100 Arezzo, Italy; Department of Biochemical Sciences “A Rossi Fanelli”, Sapienza Università di Roma, Rome, Italy

**Keywords:** Coronavirus, SARS Coronavirus, Epidemiology, Pandemics, Virus classification, BQ.1

## Abstract

The BQ.1 SARS-CoV-2 variant, also known as Cerberus, is one of the most recent Omicron descendant lineages. Compared to its direct progenitor BA.5, BQ.1 carries out some additional spike mutations in some key antigenic site which confer it further immune escape ability over other circulating lineage. In such a context, here we performed a genome-based survey aimed to obtain an as complete as possible nuance of this rapidly evolving Omicron subvariant. Genetic data suggests that BQ.1 represents an evolutionary blind background, lacking of the rapid diversification which is typical of a dangerous lineage. Indeed, the evolutionary rate of BQ.1 is very similar to that of BA.5 (7.6 × 10^−4^ and 7 × 10^−4^ subs/site/year, respectively), which is circulating by several months. Bayesian Skyline Plot reconstruction, indicates low level of genetic variability, suggesting that the peak has been reached around September 3, 2022. Structure analyses performed by comparing the properties of BQ.1 and BA.5 RBD indicated that the impact of the BQ.1 mutations on the affinity for ACE2 may be modest. Likewise, immunoinformatic analyses showed modest differences between the BQ.1 and the BA5 potential B-cells epitope. In conclusion, genetic and structural analysis on SARS-CoV-2 BQ.1 suggest that, it does not show evidence about its particular dangerous or high expansion capability. The monitoring genome-based must continue uninterrupted for a better understanding of its descendant and all other lineages.

## 1. Introduction

Nowadays, the world continues its fight against the COVID-19 pandemic declared by the World Health Organization (WHO) on March 11^th^, 2019 (https://www.who.int/director-general/speeches/detail/who-director-general-s-opening-remarks-at-the-media-briefing-on-covid-19---11-march-2020). As of November 6, 2022, the WHO stated that there were over 629 million confirmed cases and over 6.5 million deaths due to COVID-19 (https://www.who.int/publications/m/item/weekly-epidemiological-update-on-covid-199-november-2022). This pandemic has been caused by the coronavirus SARS-CoV-2, which was identified for the first time in December 2019 during a pneumonia outbreak in Wuhan (China) [1]. Along the pandemic course, SARS-CoV-2 has evolved with the generation of many new variants endowed with progressive increase in transmissibility [2], hence perpetuating the pandemia despite the availability since early 2021, of safe and effective vaccines. With time, the virus variants have shown an increasing trend to escaping from antibody immunity conferred by the vaccines (https://www.who.int/health-topics/vaccines-and-immunization#tab=tab_1), and infections are likely to remain a problem for the time being in most countries. SARS-CoV-2 is a positive-sense, single-stranded RNA virus, of which replication and transcription are made by the viral RNA-dependent RNA polymerase (RdRp, nsp12) [3], and during its RNA replication, very often errors occur [4]. Owing to this rapid evolution, mainly shaped by natural selection, the formation of new variants is not a novelty or a sporadic case, but it is a certainty that periodically recurs [2, 5]. Accordingly, many variants emerged since pandemia start in China and continue to emerge till the present days. Some of these variants have been given Greek names (i.e. Alpha, Beta, Gamma, Delta, and Omicron) and declared by the WHO to be of particular concern because of their high transmissibility and impact on populations health (https://www.who.int/news/item/31-05-2021-who-announces-simple-easy-to-say-labels-for-sars-cov-2-variants-of-interest-and-concern). Since the end of 2021, the Omicron variant has out-competed previous variants, becoming the dominant one worldwide and continuing to spread through the generation of a number of sub-variants, which include BA.1, BA.2, BA.3, BA.4, BA.5, and descendant lineages (https://www.who.int/activities/tracking-SARS-CoV-2-variants).

One of the most recent descendant lineages is represented by the Omicron subvariant BQ.1, also known as Cerberus [6]. BQ.1 is a direct descendant of BA.5 and carries out some additional spike mutations in some key antigenic site (K444T and N460K), and its first descendant, BQ.1.1 (as of November 1, 2022 there are at least 5 further descendants), carries a further additional mutation (R346T) [6]. As of October 9, 2022 BQ.1 has been detected in 65 countries with an overall prevalence of 6% (https://www.who.int/news/item/27-10-2022-tag-ve-statement-on-omicron-sublineages-bq.1-and-xbb). Considering that these additional mutations confer further immune escape ability over other circulating lineage [7-8] a constant genome-based monitoring is mandatory.

In such a context, we have here devised an approach whereby results on genetic variability/phylodynamics, structural and immunoinformatics were compared and integrated in order to obtain an as complete as possible nuance of this rapidly evolving and potentially high transmissible Omicron subvariant. More specifically, the objective of our research was to gather data that could validate or argue against the concern about a worldwide epidemiological expansion of such a strongly immune-evasive variant as the Omicron BQ.1.

## 2. Materials and Methods

First genomic epidemiology of SARS-CoV-2 BQ.1 Omicron variant (GSAID Clade 22E) has been reconstructed by using a subsampling focused globally over the past 6 months, built with nextstrain/ncov (https://github.com/nextstrain/ncov), available at https://gisaid.org/phylodynamics/global/nextstrain/, includng all genomes belonging to the GSAID Clade 21 M (Omicron). After the first genomic assessment, a subset of 1575 genome was build up including, together with all genomes of BQ.1 for which complete sampling date was available, representatives of the main variant of concern (BA.1, BA.2, BA.3, BA.4, BA.5). Genomes were aligned by using the algorithm L-INS-I implemented in Mafft 7.471 [9], producing a dataset 29,645 bp long. Manual editing was performed by using the software Unipro UGENE v.35 [10]. The software jModeltest 2.1.1 [11] was used to find the best probabilistic model of genome evolution with a maximum likelihood optimized search. Phylogenomic relationship among variants and time of divergence were investigated by using Bayesian Inference (BI), which was carried out using the software BEAST 1.10.4 [12] with runs of 200 million generations under several demographic and clock model. In order to inference on the best representative output the selection of the better model was performed by the test of Bayes Factor [13] by comparing the 2lnBF of the marginal likelihoods values following Mugosa et al. [14]. The phylogenetic trees were edited and visualized using FigTree 1.4.0 (available at http://tree.bio.ed.ac.uk/software/figtree/). The software Beast was also used to co-estimate the evolutionary rate, Bayesian Skyline Plot (BSP) and lineages thorough times for BQ.1 variant (with a subset of 1114 genomes), with runs of 300 million generations under the Bayesian Skyline Model with the uncorrelated log-normal relaxed clock model. All datasets were build by downloading genomes form GSAID portal (https://gisaid.org/) available at October 15, 2022. See Table S1 for details on the genomes included into the dataset and Authorship.

Homology models of the mutant BQ.1 Spikes were created by means of the software Modeller 10.3 [15]. Surface electrostatic potential was calculated and displayed with the graphic program PyMOL [16]. Net charge were calculated by the software PROPKA3 [17]. Foldx5 [18] was applied to optimize the side chain conformation of the models built by Modeller using the Foldx function “RepairPDB”. Interaction energy between the Spike RBD and ACE2 were predicted with the Foldx5 function “AnalyseComplex”. Interface residue-residue interactions were assessed with the Foldx5 “PrintNetwork” function. In silico mutagenesis was obtained with the built-in functions available within PyMOL. In silico alanine scanning of the residues at the interface between RBD and ACE2 was carried out using the method available via the web server DrugScore(PPI) [19], available at https://cpclab.uni-duesseldorf.de/dsppi/. The method is a fast and accurate computational approach to predict changes in the binding free energy when each residue at the subunit interface is mutated into alanine.

DiscoTope 2.0 has been used to predict discontinuous B cell epitopes from protein three dimensional structures using the default DiscoTope score Threshold of -3.700 [20].

## 3. Results

Phylogenomic reconstruction (Fig. 1) indicates that genomes of BQ.1 (GSAID Clade 22E) clustered within the not-monophyletic GSAID Clade 21L with close relationship with its direct progenitor BA.5. Results of the Bayes Factor on the dataset of 1575 genomes revealed that the Bayesian Skyline Model under the lognormal uncorrelated relaxed clock model fitted data significantly better than other tested demographic and clock models. The Maximum Clade Credibility tree (Fig. 2), indicates that all genomes of BQ.1 clustered together in a monophyletic group whose the common ancestor is temporally placed 128 days before October 10, 2022 (i.e. June 5, 2022). Bayesian Skyline Plot (BSP) (Fig. 3) showed that viral population has undergone an increase in size starting from 64 days before October 10, 2022 (i.e. August 7, 2022), reaching the peak about 37 days before October 10, 2022 (i.e. September 3, 2022). Lineages through times plot (Fig. 4) indicates that the increase of the number of lineages started about 110 days before October 10, 2022 (i.e. June 22, 2022). Evolutionary rate co-estimated with BSP and lineages through times amount to 7.6×10^−4^ [95% HPD 5.2×10^−4^ - 9.8×10^−4^] subs/sites/years.

**Figure 1.**
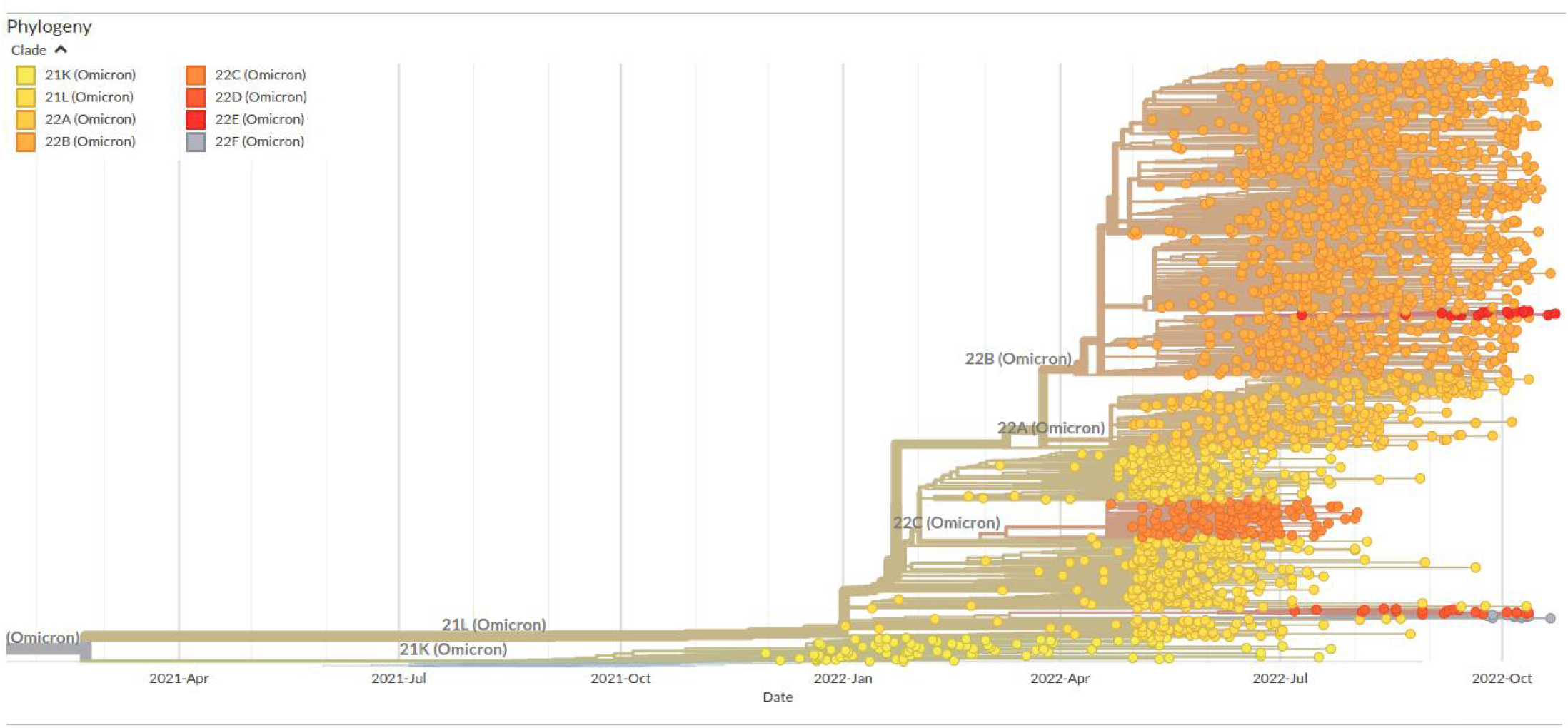
Highlight of the Omicron Clade (GSAID Clade 21L) in the time-scaled phylogenetic tree of a representative global subsample of 2809 SARS-CoV-2 genomes sampled between January 2022 and November 2022. Figures has been edited by using the software GIMP 2.8 (available at https://www.gimp.org/downloads/oldstable/).

**Figure 2.**
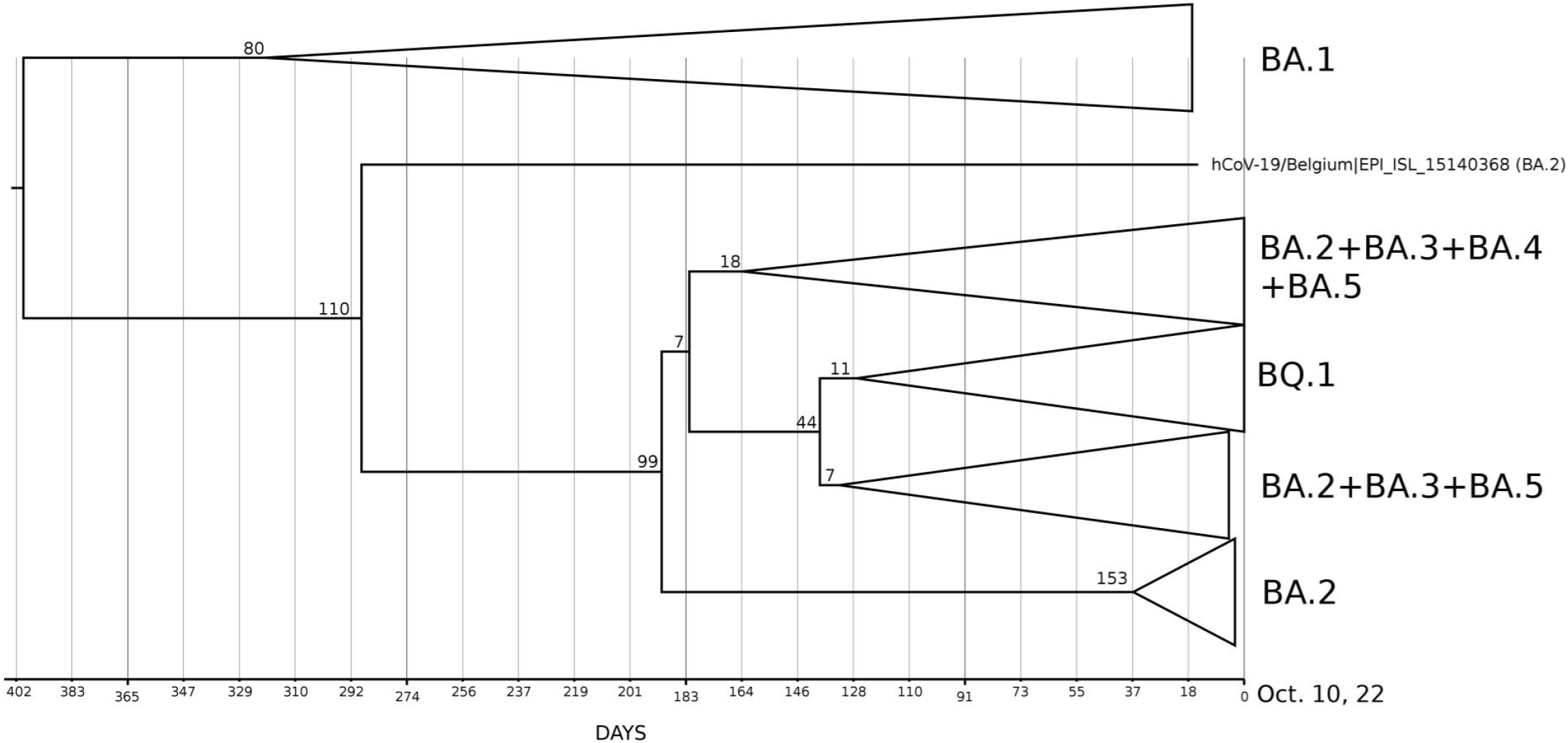
Maximum Clade Credibility Tree estimated from 1575 whole genomes of BQ.1, BA.1, BA.2, BA.3, BA.4 and BA.5 downloaded from GSAID portal (https://gisaid.org/) available at October 15, 2022. See Table S1 for details on the genomes included in the analyses. All showed nodes are well-supported. The bar under the tree indicates the time scale expressed in days before October 10, 2022 which represent the most recent sampling date included in the analyzed dataset. Node values indicates branches time expressed in days.

**Figure 3.**
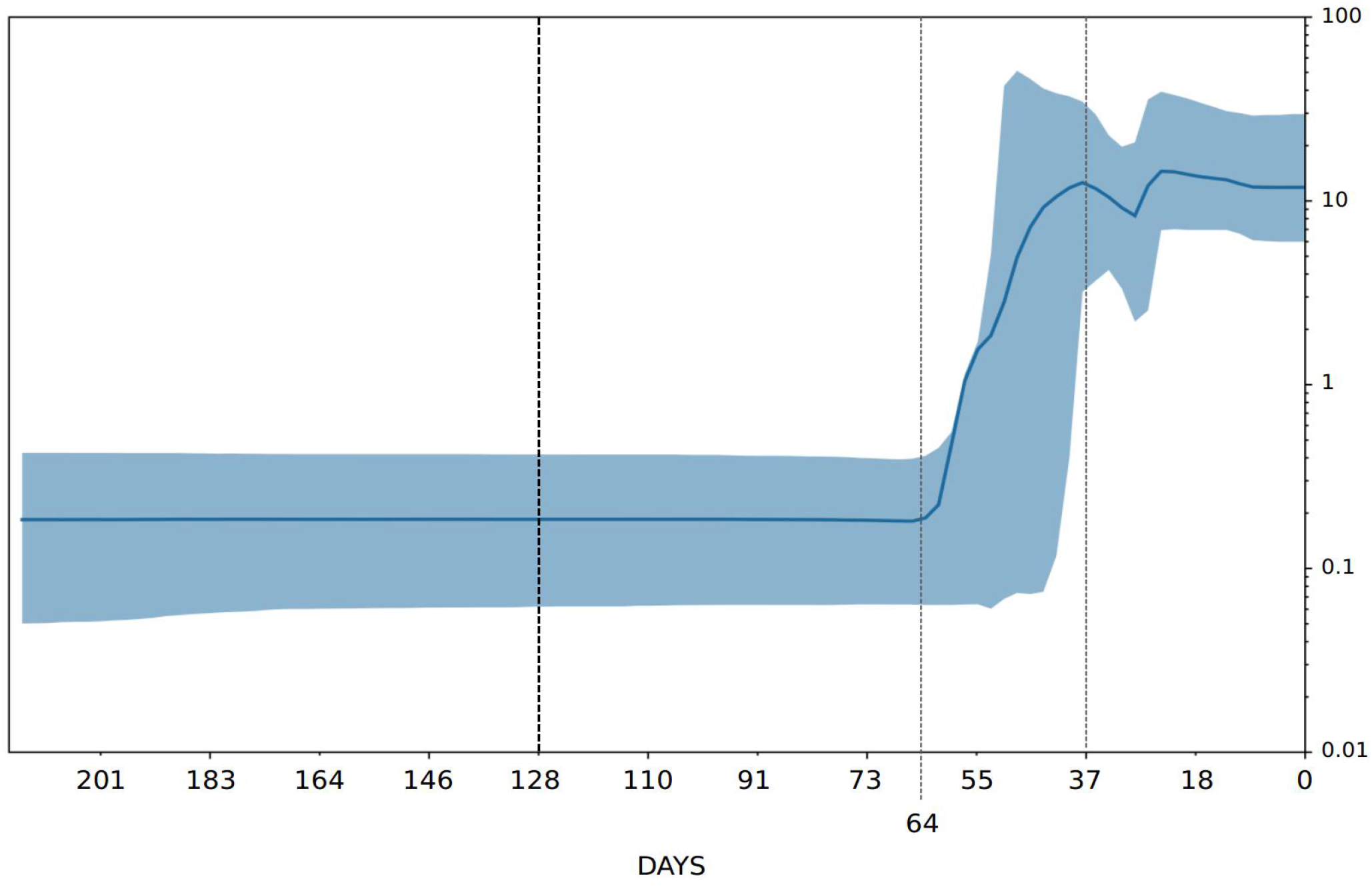
Bayesian skyline plot of SARS-CoV-2 BQ.1 variant. The viral effective population size (y-axis) is shown as a function of days (x-axis). B). SARS-CoV-2 BA.2.75 variant lineages through time. Number of lineages (y-axis) is shown as a function of days (x-axis). Solid area represents the 95% high posterior density (HPD) region.

**Figure 4.**
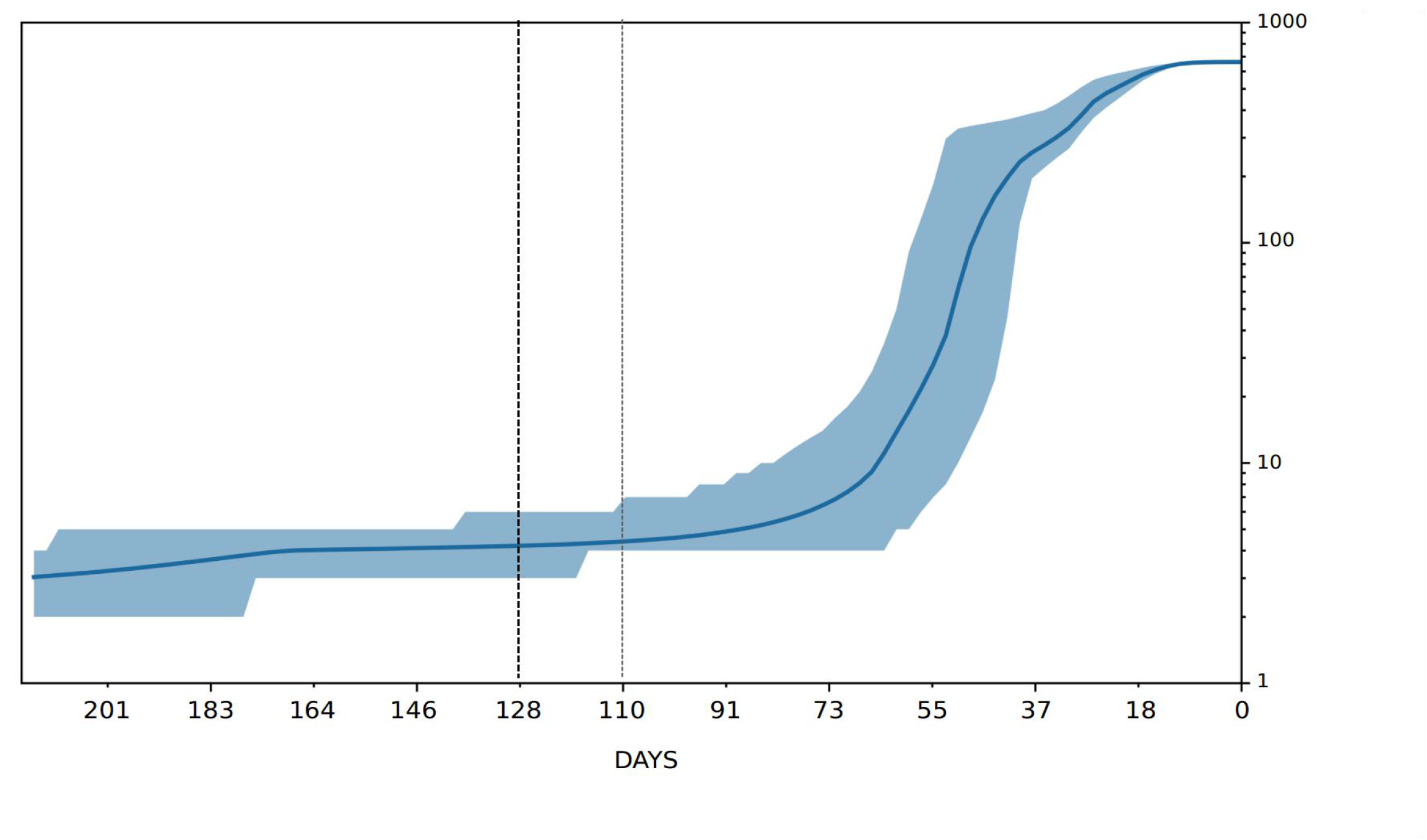
SARS-CoV-2 BQ.1 variant lineages through time. Number of lineages (y-axis) is shown as a function of days (x-axis). Solid area represents the 95% high posterior density (HPD) region.

The predicted structural properties of the BQ.1 Spike have been compared with those of BA.5 Spike. The characterizing BQ.1 mutations are reported in Table S2. The N-terminal domains (NTD) of the two Spikes are identical while the Receptor Binding Domains differ for the three BQ.1 mutations R346T, K444T and N460K. N460K has been reported also in BA.2.75. Homology modelling of the BQ.1 Spike indicates that the R346T mutation occurs in a loop exposed to the solvent while K444T is inside the Receptor Binding Motif near the interface to ACE2 (Fig. 5). N460K occurs also in an exposed loop not directly involved in the interaction with ACE2 (Fig. 5). The predicted net charge of the RBD at pH=7.0 (set as a reference pH, though not necessarily reflecting the physiological environment) was calculated by means of the program PROPKA3. The predicted overall net charge is dependent on the combination of the sidechain charges. The net charge is influenced by the interactions of the side chains with the surrounding atoms. To sample the variability of the interactions, 100 homology models of each RBD have been calculated with Modeler. Indeed, the Modeler refinement stage of the homology modelling can produce models differing for the conformation of side chains. Each model has been optimized by the Foldx5 “RepairPDB” procedure and the net charge has been calculated by PROPKA3. The final charge was the average of the 100 charges and the variability estimated by the standard error at 95% confidence interval. The same procedure was applied to estimate the interaction energy of the complex ACE2-RBD. The net charge attributed to the BA.5 and BQ.1 identical NTD Spike moieties is +1.18 ± 0.09. The BA.5 and BQ.1 RBD shows a net charge equal to +5.18 ± 0.03 and +4.20 ± 0.03, respectively. The decreased charge is coherent with the replacement of the two positively charged residues R346 and K444 by the uncharged polar Thr. The pattern is confirmed by the comparison of the electrostatic potential surfaces of the two RBDs. The surface of the BA.5 RBD has a more positively charged region with respect to the corresponding in BQ.1 (see Fig. S1). According to visual and Foldx5 interface analyses, Thr444 does not directly interact with ACE2 (Fig. 5). Interaction energies between ACE2 and BA.5 and BQ.1 estimated by Foldx5 are -4.67 ± 0.55 and -4.50 ± 0.57 Kcal/mol, respectively. Apparently, the interaction between ACE2 and BQ.1 is predicted to be slightly weaker than in the case of BA.5 within the limits of the method. Even if the influence of the characterizing mutations of BQ.1 RBD appear marginal on the interaction with ACE2, they may have an impact on the interaction with antibodies. As an example, the interaction with the monoclonal antibody CV38-192 reported in the PDB data set 7LM8 has been examined. The RBD positions 346 and 444 are at the interface with the antibody heavy chain (Fig. 5). Foldx5 analysis suggests that in the wild type RBD the residue R346 interacts with E54 and Y52 while K444 interacts with D56 of the heavy chain. Removal of the positive charges of R346 and K444 should disrupt the salt bridges. In silico mutagenesis have been applied to replace the single and both positions by Threonine, as in BQ.1. Interaction energies have been predicted by Foldx5. The results suggest that the mutations tend to destabilize the interaction with the antibody heavy chain (Table 1). Moreover, the analysis by DrugScore (PPI) confirms that R346 is a major hotspot. Indeed, the loss of interaction energy of the complex upon mutation of R346 to alanine is rather high and equal to G=1.82 Kcal/mol.

**Figure 5.**
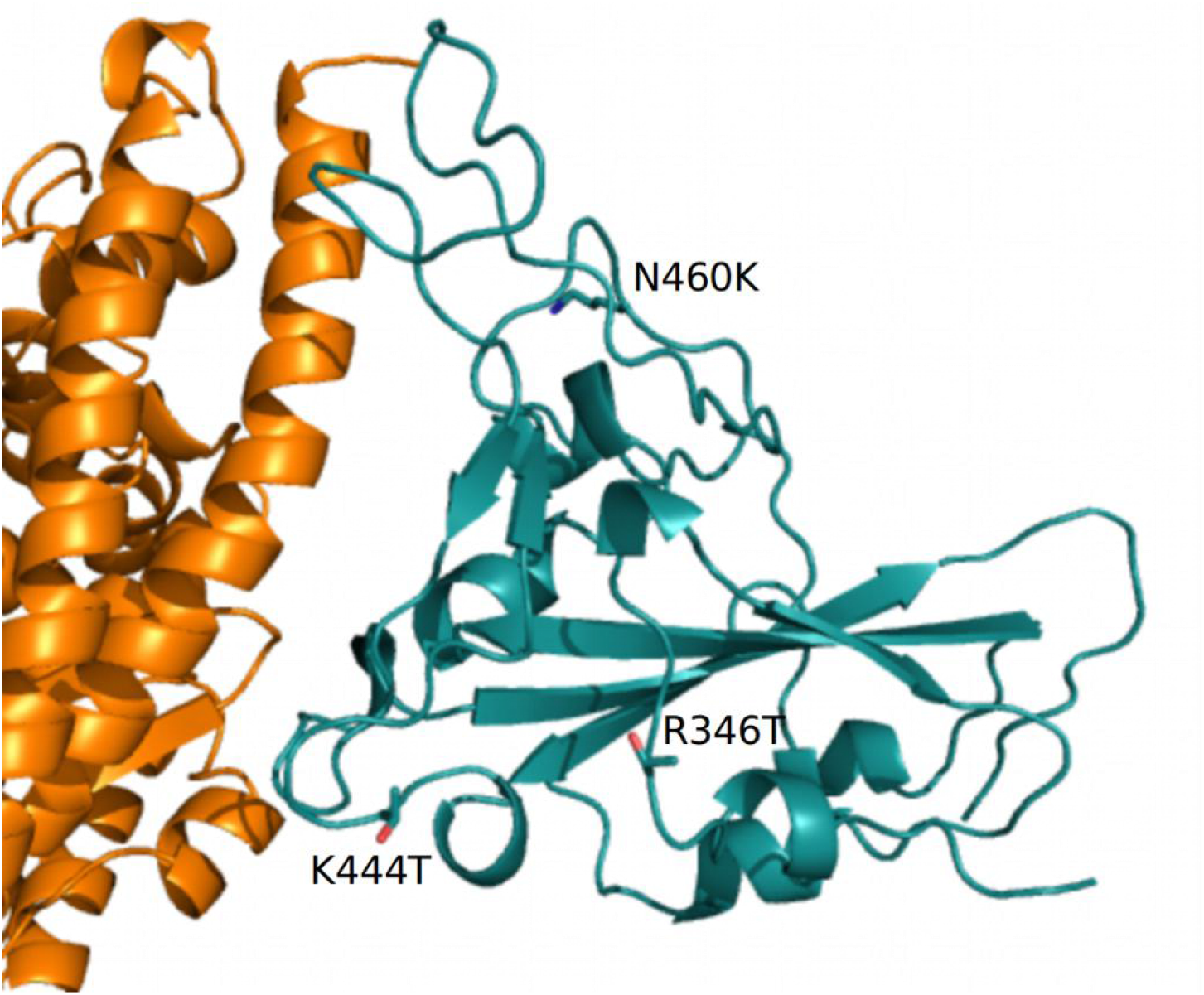
Complex between ACE2 (orange) and RBD (deep-teal) represented as cartoon models. The side chains of the residue mutations specific to BQ.1 are displayed with stick models and are labeled. Only part of ACE2 receptor is displayed.

**Figure 6.**
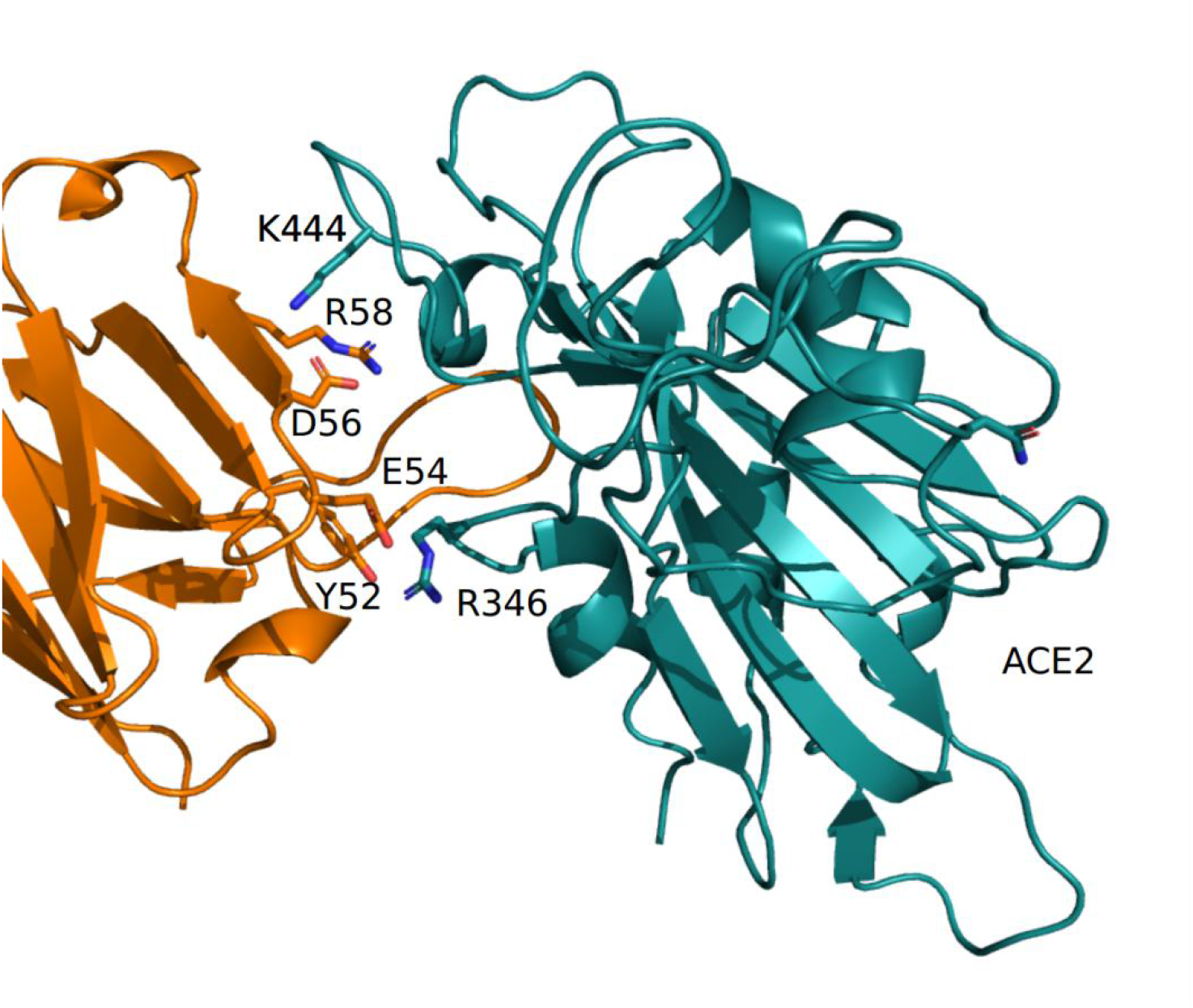
Complex between the COV38-142 antibody heavy chain (orange) and RBD (deep-teal) as reported in the PDB coordinate set 7LM8. Interacting side chains are displayed with stick models and are labeled. The label «ACE2» marks the interface to the receptor.

**Table 1.**
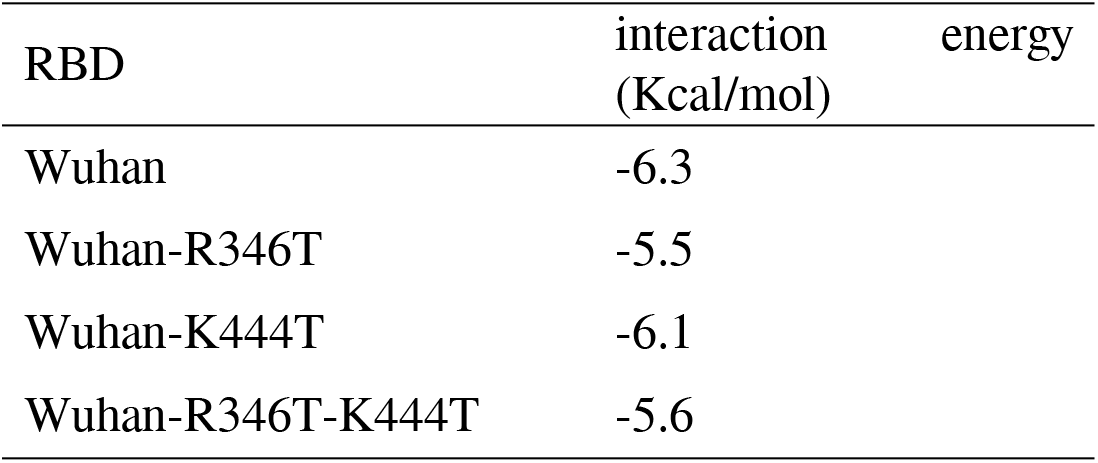
Interaction energy between Wuhan and mutant RBDs and the CV38-142 antibody heavy chain calculated by Foldx5

The immunoinformatic analysis has found the presence of 108 B-cells epitope residues in the BQ.1 strain and 111 B-cells epitope residues in the BA.5 strain of the K439L mutation of BQ.1. The main differences between BA.5 and BQ.1 have been reported in Table 2.

**Table 2.** Defining Spike glycoprotein aminoacidic mutations for BA5 and BQ1.1 SARS-CoV-2 variants with the respective discotope predicating scores.

| BA5 | Amino acidic position | BQ.1 | BQ.1 DiscoTope score | BA5 DiscoTope score |
| --- | --- | --- | --- | --- |
| SER | 438 | SER | -3.733* | -2.003 |
| LYS | 439 | THR | 0.220 | 1.904 |
| VAL | 440 | VAL | 3.112 | 4.307 |
| GLY | 441 | GLY | 2.203 | 3.670 |
| GLY | 442 | GLY | 0.553 | 2.047 |
| ASN | 443 | ASN | -3.125 | -1.276 |
| TYR | 444 | TYR | -1.383 | 0.031 |
| ASN | 445 | ASN | -2.964 | -1.719 |
| LEU | 487 | LEU | -4.250* | -3.559 |
| GLN | 488 | GLN | -4.265* | -3.436 |
| SER | 489 | SER | -3.458 | -2.349 |
| TYR | 490 | TYR | -4.915* | -3.658 |
| GLY | 491 | GLY | -0.377 | 0.794 |
| PHE | 492 | PHE | -3.202 | -1.742 |
| ARG | 493 | ARG | 0.628 | 2.089 |
| PRO | 494 | PRO | 1.352 | 2.664 |
| THR | 495 | THR | 2.953 | 3.742 |
| TYR | 496 | TYR | -0.290 | 0.704 |
| GLY | 497 | GLY | 1.939 | 2.529 |
| VAL | 498 | VAL | -1.746 | -0.966 |
| GLY | 499 | GLY | -2.855 | -2.103 |
| HIS | 500 | HIS | -2.180 | -1.223 |
| GLN | 501 | GLN | -3.523 | -2.327 |

## 4. Discussion

The SARS-CoV Omicron BQ.1 variant represents one of the most recent discovered lineages and require an in-depth study on its capacity for expansion and contagiousness. Here we sought for a deep insight into the evolutionary and structural patterns of the SARS-CoV BQ.1 variant by means of all genomes available in GSAID at October 15, 2022.

Phylogenomic reconstruction indicates that genomes of BQ.1 (GSAID Clade 22E) clustered within the wide GSAID Clade 22B (BA.5). This is not surprising considering that BQ.1 is a descendant of BA.5. One of the most striking result is given by the evolutionary condition of BQ.1, indeed, phylogeny suggested an evolutionary condition of BQ.1 very similar to the variants BA.2.75 (GSAID Clade 22D) and BA.2.12.1 (GSAID Clade 22C), which represent an evolutionary blind background with no further epidemiologically relevant descendant. If this evolutionary condition is confirmed in the near future, it will be possible to appreciate in BQ.1 and its descendant an atypical accumulation of neutral loss-of-function mutations. Indeed, typically these kinds of lineages are characterized by several nucleotidic mutations with no (or very few) ammino acidic mutations for what important genes are concerned. In addition, the branches’ length in the phylogenetic tree suggested the lack of a rapid diversification, which is typical of a dangerous lineage at the beginning of its evolutionary path.

Phylodynamic reconstruction performed on a dataset comprising 1575 genomes, indicates that among the included variants (BQ.1, BA.1, BA.2, BA.3, BA.4, BA.5 and their descendant sublineages) BA.1 and BQ.1 are the only ones to show a monophyletic condition. In the Maximum Clade Credibility tree the common ancestor to all genomes of BQ.1 is temporally placed 127 days before October 10, 2022 (which is the most recent collection date), i.e. June 5, 2022. This dating predates of about one month the first detected genome of BQ.1 (for which a complete sampling date is available), which was isolated in Nigeria (Abuja) on July 4 and initially labeled as BA.5-like (and then relabeled as BQ.1). It is interesting to note that considering its date of origin, the virus circulated undisturbed for a long time before being detected. This is not the feature of a dangerous variant, which typically explodes much faster in terms of numbers of infections and population size. Accordingly, Bayesian Skyline Plot (BSP) reconstruction, estimated on 1114 whole genomes of BQ.1 (collection range 2022/07/04 - 2022/10/10), indicates low level of genetic variability, also confirmed by genetic distance matrix, where the largest value amounts to 0.731 (±0.002) and has been found in few cases between genomes sampled three months later. Indeed, after an initial low and flattened level of genetic variability, around August 7, 2022 (64 days before October 10, 2022) the expansion of the viral population size started with a very steep curve that lasted for 18 days, after which the peak has been reached around September 3, 2022. The reconstruction of lineages through time indicates that the increasing of the number of lineages started about 10 days before the beginning of the increasing of the population size, as usual in this kind of cases. During the plateau phase genetic variability (and accordingly viral population size) have fluctuated with some rise and fall but nothing relevant, and currently viral population size appears to be stable and flattened. It should be pointed that this is not the typical trend of a lineage that is about to explode in terms of population size and accordingly, in terms of contagiousness as shown at the beginning of the pandemic when variability increased very quick with a very vertical curve (see *i*.*a*., Lai et al. [21]). On the contrary this trend is very similar to what shown for the BA.2.75 variant which similarly to BQ.1, at the beginning has aroused much concern, but after a deep genome-based survey it did not show evidence of a particular dangerous or high expansion capability being even more slow than others [22]. Indeed, BQ.1 situation is coherent with a scenario quite typical of an evolutionary lineage that presents new features in comparison to its direct progenitor (BA.5) but these new features, at the present stage, do not represent a further boost able to promote an abnormal expansion. In addition, the lack of increasing of lineages through time further confirms the lack of an enlarge in the number of haplotypes in recent times. Evolutionary rate estimated for BQ.1 amount to 7.6 × 10^−4^ subs/site/year with a very close range (5.2 × 10^−4^ - 9.8 × 10^−4^ subs/site/year). It is a further confirm of the low level of genetic variation and poor capability of demographic expansion, indeed, it is very similar to variant BA.5 with 7 × 10^−4^ subs/site/year and BA.2.75 with 1.6 × 10^−4^ subs/site/year [22]. For what the comparison with BA.5 is concerned, it should be pointed that this last one has been circulating for several months and its current level of variability, of course, is lower than in the early stages. Accordingly, the evolutionary rate of BQ.1 should be greater if it were dangerous with a highly contagious capability lineage. Indeed, at the beginning of the current pandemic, the evolutionary rate of the first lineage of SARS-CoV-2 was about 6.58e^-3^ subs/site/year [23], that means that also in this case the new variant presents a factor of 10^−1^ more slow than Wuhan-Hu-1 variant.

The comparison of the properties of BQ.1 and BA.5 RBD predicted that the impact of the BQ.1 mutations on the affinity for ACE2 may be modest although it appears to be potentially destabilizing. In fact, the calculation of the RBD net charge, that is an indirect measure of the dominant charge of the electrostatic potential surface, suggests that it is decreased in BQ.1. The value of the RBD net charge in BQ.1 is comparable to that the Delta variant and would suggest a diminution rather than an increase of virus transmissibility compared to the parental Omicron BA.5 variant [24]. Interestingly, two of the characterizing mutations that remove the charged residues in the Wuhan virus, R346 and K444, have a clear destabilizing effect on the interaction between RBD and the class of antibodies that recognizes the epitope containing the two positions. It may be speculated that the current evolution of the virus is optimizing its immune escape ability rather than its affinity for the receptor. The limited immunoinformatic analysis here performed has shown differences between the BQ.1 and the BA5 potential B-cells epitope. Although such difference may appear to be modest, it adds to the already established mutations of BA.5 spike epitopes in explaining the elevated capability of the BQ.1 variant to escape neutralizing antibodies generated by vaccination or infection [8]. Besides antibody responses, particularly those neutralizing virus entry into host cells, COVID-19 disease is under control of T cell-mediated immunity, particularly CD4+T and CD8+T lymphocytes, of which CD8+T ones are endowed with potent cytotoxicity against SARS-CoV-2-infected cells [25]. In this immunological scenario, it is also due to the observation that the BQ1 R346T mutation may affect a T cell recognized-immunogenic peptide that is largely preserved in all previous variants [26]. However, it is unknown whether or the extent to which arginine substitution by threonine may affect peptide immunogenicity. In addition, we have observed that all other highly preserved T cell responsive to epitopes of the Spike protein, as reported by De la Fuente et al. [26] appear to be unmodified in the BQ.1 variant. Although further, direct research on this topic is warranted, altogether, these data do not support any relevant immune evasion by BQ1 of the T cell response against the above spike epitopes.

In accordance to all evolutionary theories, often new SARS-CoV-2 variants exhibit antibodies escape capabilities [27, 28], but this is not necessarily strictly related to a high diffusion capability or to an improved pathogenicity of the virus [22]. Accordingly to the data currently available, SARS-CoV-2 BQ.1 appear as a new variant with no enhanced capabilities of infectivity or pathogenicity with respect to its direct progenitor BA.5.

## Conclusions

In conclusion, genetic and structural analysis on SARS-CoV-2 BQ.1 suggest that, although this new variant presents several spike mutations of interest and an overall highly immune-evasive of neutralizing antibodies [8], currently it does not show evidence about its particular dangerous or high expansion capability. The Omicron variant of concern remains the dominant variant circulating globally (https://www.who.int/publications/m/item/weekly-epidemiological-update-on-covid-19---9-november-2022) but BQ.1 appears even more slowly than the last variant that became dominant at the commencement of the year, the BA.5 one, and based on current data, it does not appear to present an alarming situation. However, this condition must not be understood as a reason to let down the guard against the pandemic or the generation of further variants. Indeed, new further mutations can make BQ.1 more dangerous or generate new subvariants. As of November 8, 2022 BQ.1 presents four known sublineages whose plausibly can be able to exhibits similar antibodies escape capabilities. Accordingly, the constant genome-based surveys remain the best tool for a better understanding of the phenomenon. Monitoring of BQ.1 and its descendant, as well as the monitoring of all other lineages, must continue uninterrupted in order to identify and/or predict important changes in genomic composition and/or diffusion capability.

## Acknowledgments

This research was funded from by FONDAZIONE DI SARDEGNA <u>bando 2022-2023</u> for the Dipartimento di Scienze Biomediche - UNISS (to Daria Sanna and Fabio Scarpa). Marta Giovanetti is funded by PON “Ricerca e Innovazione” 2014-2020. Authors are grateful to Cristina Giuliano for providing the list of complexes between Spike RBD and antibodies involving the sites 346 and 444. SP is in part supported by the Sapienza grant n. RP12117A7670A1E8.

We also would like to thank all the authors who have kindly deposited and shared genomes on GSAID.

## Data Availability Statement

Genomes analysed in the present study were taken from GSAID database and are available at https://gisaid.org/.

## Author Contributions

Conceptualization, F.S., D.S., M.C. (Marco Casu), P.L.F., A.C. (Arnaldo Caruso), A.C. (Antonio Cassone), S.P. (Stefano Pascarella) and M.C. (Massimo Ciccozzi); data analyses, F.S., D.B. and S.P; validation, F.S., D.S., M.C. (Marco Casu), P.L.F., D.B., S.P. (Stefano Pascarella) and M.C. (Massimo Ciccozzi); writing—original draft preparation, F.S., D.S., and M.C. (Marco Casu); writing—review and editing, F.S., D.S., D.B., A.B., I.A., M.C., P.L.F., M.G., A.M., G.C., A.C. (Arnaldo Caruso), F.C., R.C., A.C. (Antonio Cassone), S.P. and M.C. (Massimo Ciccozzi).

All authors have read and agreed to the published version of the manuscript.

## FIGURES CAPTION

**Figure S1.**
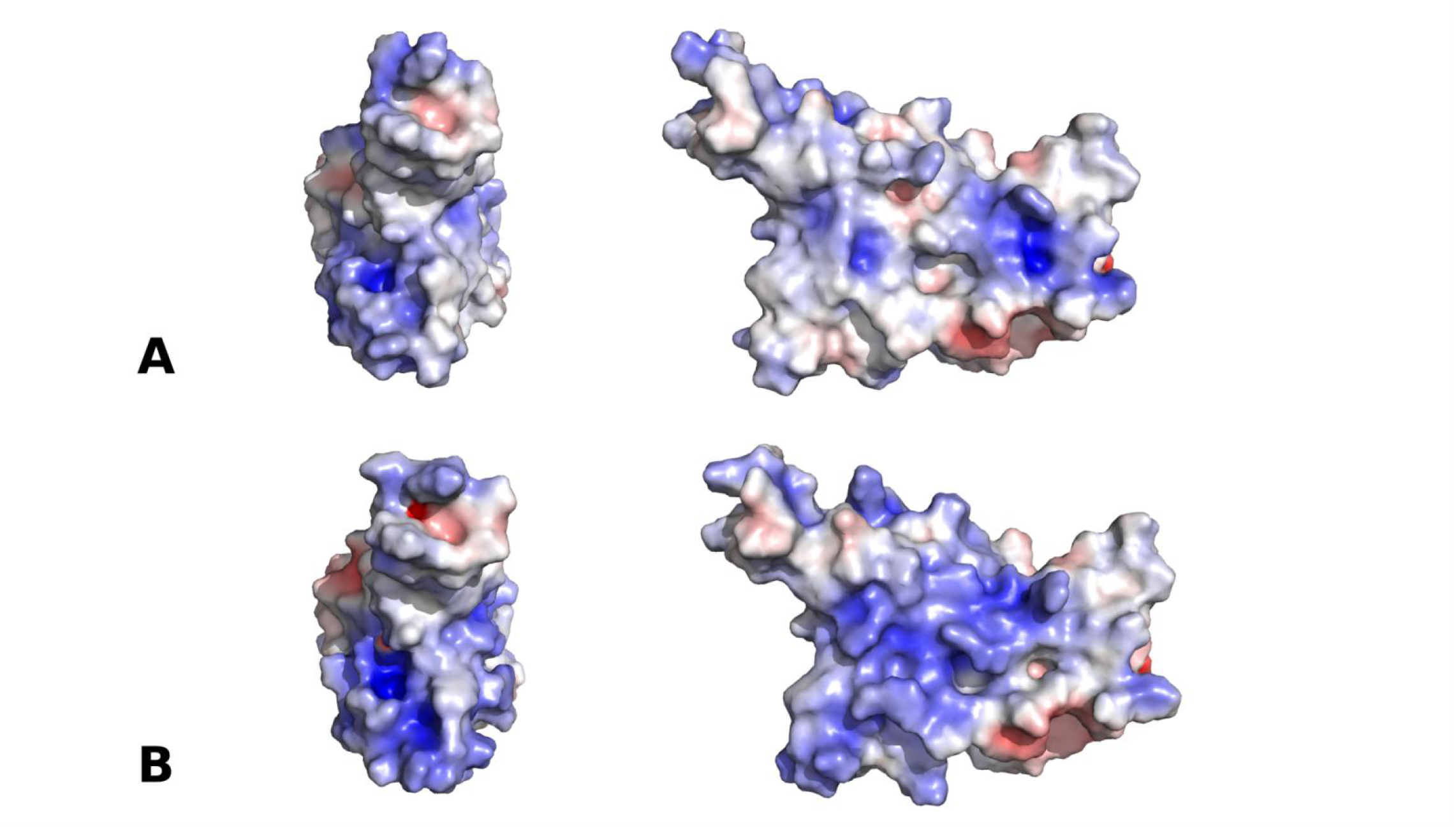
Comparison of the electrostatic potential surface of the BQ.1 (A) and BA.5 (B) RBDs. The domains are oriented with the ACE2 interface in the front (left) and rotated by 90 degrees around f the y vertical axis (right). The color scale ranges from -5.0 (dark red) to +5.0 (dark blue) kT/e

**Supplementary Table 1.**
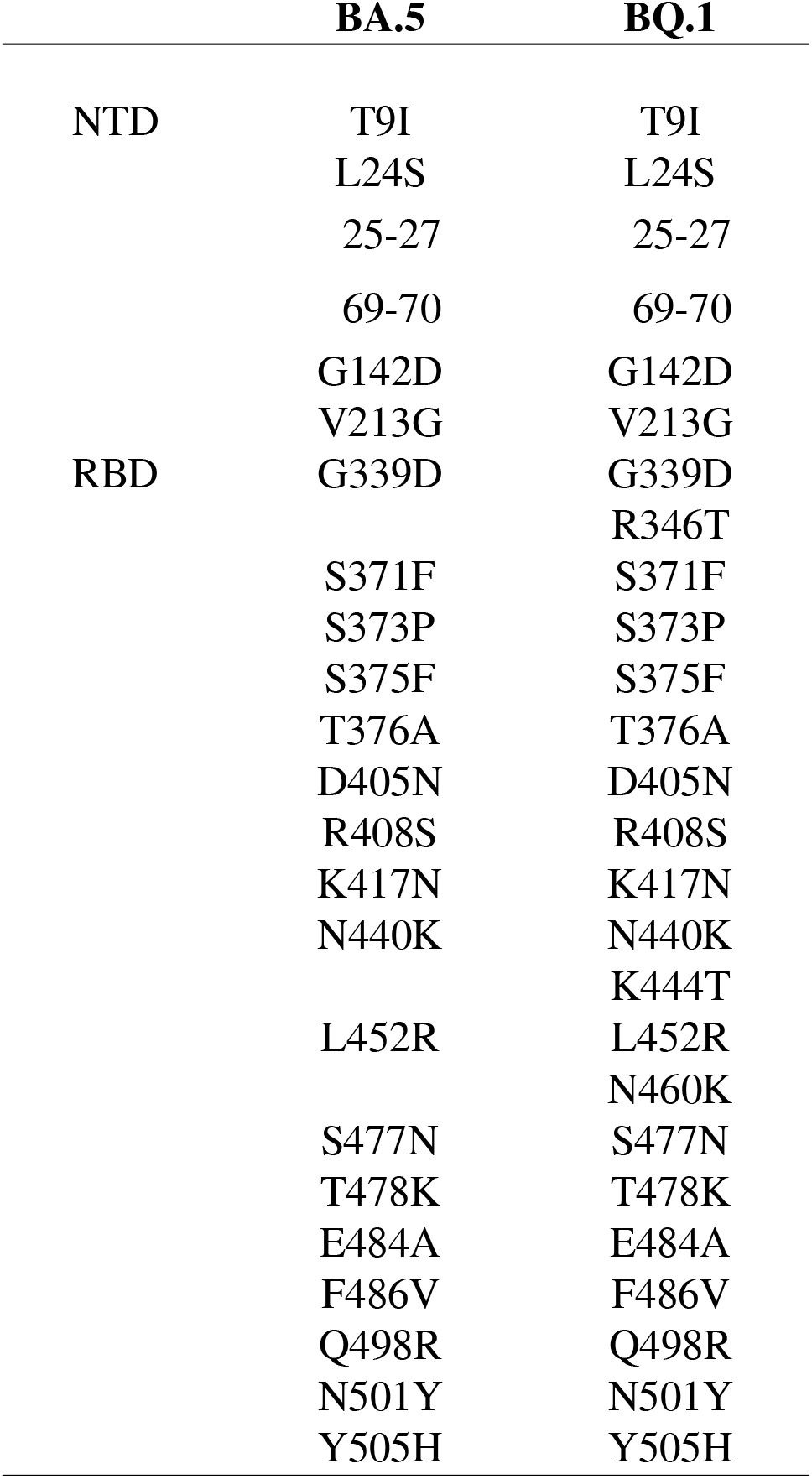
Comparison of the BA.5 and BQ.1 mutations in the NTD and RBD regions of the Spike

