## Supplementary Table 1 for "Genetic and structural data on the SARS-CoV-2 Omicron BQ.1 variant reveal its low potential for epidemiological expansion"

### SUPPLEMENTAL TABLE

**Data Availability**

GISAID Identifier: EPI_SET_221108qe
doi: 10.55876/gis8.221108qe

All genome sequences and associated metadata in this dataset are published in GISAID's EpiCoV database. To view the contributors of each individual sequence with details such as accession number, Virus name, Collection date, Originating Lab and Submitting Lab and the list of Authors, visit 10.55876/gis8.221108qe

**Data Snapshot**

- EPI_SET_221108qe is composed of 1,114 individual genome sequences.
- The collection dates range from 2022-07-10 to 2022-10-05;
- Data were collected in 35 countries and territories;
- All sequences in this dataset are compared relative to hCoV-19/Wuhan/WIV04/2019 (WIV04), the official reference sequence employed by GISAID (EPI_ISL_402124). Learn more at https://gisaid.org/WIV04.

### SUPPLEMENTAL TABLE

**Data Availability**

GISAID Identifier: EPI_SET_221109dq
doi: 10.55876/gis8.221109dq

All genome sequences and associated metadata in this dataset are published in GISAID's EpiCoV database. To view the contributors of each individual sequence with details such as accession number, Virus name, Collection date, Originating Lab and Submitting Lab and the list of Authors, visit 10.55876/gis8.221109dq

**Data Snapshot**

- EPI_SET_221109dq is composed of 91 individual genome sequences.
- The collection dates range from 2022-07-01 to 2022-09-19;
- Data were collected in 25 countries and territories;
- All sequences in this dataset are compared relative to hCoV-19/Wuhan/WIV04/2019 (WIV04), the official reference sequence employed by GISAID (EPI_ISL_402124). Learn more at https://gisaid.org/WIV04.

### SUPPLEMENTAL TABLE

**Data Availability**

| | |
|---|---|
| GISAID Identifier: | EPI_SET_221109fb |
| doi: | 10.55876/gis8.221109fb |

All genome sequences and associated metadata in this dataset are published in GISAID's EpiCoV database. To view the contributors of each individual sequence with details such as accession number, Virus name, Collection date, Originating Lab and Submitting Lab and the list of Authors, visit 10.55876/gis8.221109fb

**Data Snapshot**

- EPI_SET_221109fb is composed of 92 individual genome sequences.
- The collection dates range from 2022-08-21 to 2022-10-07;
- Data were collected in 22 countries and territories;
- All sequences in this dataset are compared relative to hCoV-19/Wuhan/WIV04/2019 (WIV04), the official reference sequence employed by GISAID (EPI_ISL_402124). Learn more at https://gisaid.org/WIV04.

### SUPPLEMENTAL TABLE

**Data Availability**

| | |
|---|---|
| GISAID Identifier: | EPI_SET_221109kc |
| doi: | 10.55876/gis8.221109kc |

All genome sequences and associated metadata in this dataset are published in GISAID's EpiCoV database. To view the contributors of each individual sequence with details such as accession number, Virus name, Collection date, Originating Lab and Submitting Lab and the list of Authors, visit 10.55876/gis8.221109kc

**Data Snapshot**

- EPI_SET_221109kc is composed of 94 individual genome sequences.
- The collection dates range from 2022-08-09 to 2022-09-26;
- Data were collected in 8 countries and territories;
- All sequences in this dataset are compared relative to hCoV-19/Wuhan/WIV04/2019 (WIV04), the official reference sequence employed by GISAID (EPI_ISL_402124). Learn more at https://gisaid.org/WIV04.

### SUPPLEMENTAL TABLE

**Data Availability**

| | |
|---|---|
| GISAID Identifier: | EPI_SET_221109vo |
| doi: | 10.55876/gis8.221109vo |

All genome sequences and associated metadata in this dataset are published in GISAID's EpiCoV database. To view the contributors of each individual sequence with details such as accession number, Virus name, Collection date, Originating Lab and Submitting Lab and the list of Authors, visit 10.55876/gis8.221109vo

**Data Snapshot**

- EPI_SET_221109vo is composed of 98 individual genome sequences.
- The collection dates range from 2022-07-21 to 2022-09-29;
- Data were collected in 16 countries and territories;
- All sequences in this dataset are compared relative to hCoV-19/Wuhan/WIV04/2019 (WIV04), the official reference sequence employed by GISAID (EPI_ISL_402124). Learn more at https://gisaid.org/WIV04.

### SUPPLEMENTAL TABLE

#### Data Availability

| | |
|---|---|
| GISAID Identifier: | EPI_SET_221109om |
| doi: | 10.55876/gis8.221109om |

All genome sequences and associated metadata in this dataset are published in GISAID's EpiCoV database. To view the contributors of each individual sequence with details such as accession number, Virus name, Collection date, Originating Lab and Submitting Lab and the list of Authors, visit 10.55876/gis8.221109om

#### Data Snapshot

- EPI_SET_221109om is composed of 86 individual genome sequences.
- The collection dates range from 2022-09-09 to 2022-10-05;
- Data were collected in 17 countries and territories;
- All sequences in this dataset are compared relative to hCoV-19/Wuhan/WIV04/2019 (WIV04), the official reference sequence employed by GISAID (EPI_ISL_402124). Learn more at https://gisaid.org/WIV04.
